# NeuroNella: A Robust Unsupervised Algorithm for Identification of Neural Activity from Multielectrode Arrays

**DOI:** 10.1101/2024.06.05.597557

**Authors:** C. Germer, D. Farina, S.N. Baker, A. Del Vecchio

## Abstract

We introduce NeuroNella, an automated algorithm developed for the identification of neuronal activity from multichannel electrode arrays. In evaluations conducted on recordings from implanted probes in the nervous system of rodents and primates, the algorithm demonstrated remarkable accuracy, showcasing an error rate of less than 1% compared to ground-truth patch clamp signals. Notably, the proposed algorithm handles large datasets efficiently without the necessity of a GPU system. The results highlighted the algorithm’s efficacy in detecting sources in a wide amplitude range and its adaptability in accommodating minor probe shifts. Moreover, the high robustness exhibited by the algorithm in decomposing recordings lasting up to 30 minutes underscores its potential for enabling longitudinal studies and prolonged recording sessions, thus opening new avenues for future brain/machine interface applications.

## INTRODUCTION

Accurate transmission of information allows biological and non-biological systems to interact and communicate with the environment. Communication between and within biological structures is due to the coordinated activity of neural networks in the brain and spinal cord. This communication takes place at the level of individual neurons and populations of neurons, and can be investigated from the firing activity of single neural cells discriminated from electrophysiological techniques^1^.

Classic invasive electrode systems placed in neural tissues maximize the selectivity of each electrode, i.e., the signal-to-noise ratio of electric signals of a few neurons is maximized with respect to the surrounding neuronal activities^2–5^. The advent of novel technologies, such as CMOS sensors^6,7^, allows the fabrication of thousands of electrodes with very close spacings; each electrode can record action potentials from multiple neural sources. Because of the increase in signal complexity, more complex algorithms with respect to classic clustering methods have been proposed^8^ to identify these neural ensembles. Yet, most of the current software is still based on conventional spike sorting approaches^9–12^. Spike sorting algorithms are classically based on segmentation and classification^11–14^, which assumes that the probability of action potentials overlapping over time is relatively low. When action potentials overlap in the same time interval, however, the identification of their waveforms and therefore the association to specific neurons is challenging. Overlapping of action potentials rapidly increases with decreasing selectivity. A theoretical approximation^15^ indicates that the number of overlapped action potentials per unit time *Ov* depends on the number of active detected neurons *N*, their average firing frequency *f*, and the action potential waveform duration *d*, according to the relation *Ov=f*^*2*^*·d·N*. As the number of detected neurons increases, the overlap rate increases linearly. For example, for 2 detected neurons, with an average firing at 50 Hz, and action potential duration of 1 ms, the average overlap rate would be only 5 action potentials/s, but when the number of detected neurons slightly increases to 10 the overlap rate increases to 25 action potentials/s. Therefore, overlapping of action potentials in time becomes a major challenge as soon as the selectivity of each electrode is slightly reduced. Conventional spike sorting approaches that segment the recording in time intervals containing action potential waveforms and cluster the segmented waveforms, without resolving action potentials overlapped in time, can only be applied with very selective electrodes. Attempts to identify overlapped potentials require complex calculations; these may not be feasible if there is a need for online sorting.

CMOS sensor technology opened up a new way of extracting neural data due to the high density of the electrodes and the fact that each electrode records multiple sources. Because of the rich information recorded by these matrices, conventional spike sorting fails in providing accurate identification of all neural firings. While new methods based on source separation have been proposed for neural decoding^16^, none of the current methods leverages the main characteristics of the neuronal sources, i.e., their *spatiotemporal sparsity*. Temporal sparsity is a strong prior that is valid for any recording from neural (and muscular) tissues^17,18^. It implies that the sources to be identified can be modelled by Dirac delta functions with a certain probability of occurrence over time (temporal sparsity) and a local distribution in space (spatial sparsity).

Here we propose a new blind source separation approach for decoding neural signals which is solely based on the assumption that the sources to be decoded are sparse in both time and space. The method therefore can separate neural activity without being influenced by the action potential overlap rate and without any further a-priori assumptions on the statistics of neural activity. We demonstrate that a blind iterative update procedure that maximizes the sparsity of each estimated source allows the identification of action potential firing times, as long as each action potential waveform is unique. Importantly, the method was extensively validated on data from concurrent recordings of neuronal action potentials by patch-clamp and multielectrode arrays. We extracted data from different anatomical structures ranging from the mouse retina to the brainstem of anesthetized macaques. Our results show that the proposed method is highly reliable in providing robust results and does not mix action potentials from nearby sources, even in cases of overlapping of action potentials. Finally, our new method works in a fully automatic way.

## RESULTS

### Decomposition algorithm

NeuroNella is a fully automatic algorithm that blindly identifies sources from multielectrode extracellular recordings. Instead of classification based on the waveform shape, NeuroNella separates action potentials by spatio-temporal filters that maximize the sparsity of the estimated sources (see Methods for details). Due to the confinement of the neuron activity in a few neighbouring electrodes (spatial sparsity), the algorithm starts by spatial segmentation of the dense probe into smaller sub-groups of channels to reduce the computational burden and to increase the probability of identification of sources with small signal amplitude. This classification algorithm clusters the channels into sub-groups with N desired channels, taking into account the signal energy (root-mean-square) and location in the probe. It therefore provides sub-groups with similar number of channels, but with different geometry (Fig. 1, see Methods for details). Each sub-group of neighbouring electrodes contains the spatial distribution of one or more neuronal activities. Segments of 25 channels have provided a good trade-off of total processing time and number of identified sources in our tests, although this parameter can be varied depending on the application. The algorithm proceeds to the decomposition of some or all spatial segments of the probe, starting with sub-groups with higher energy, as this indicates those most likely to contain neuronal activity.

**Fig. 1.**
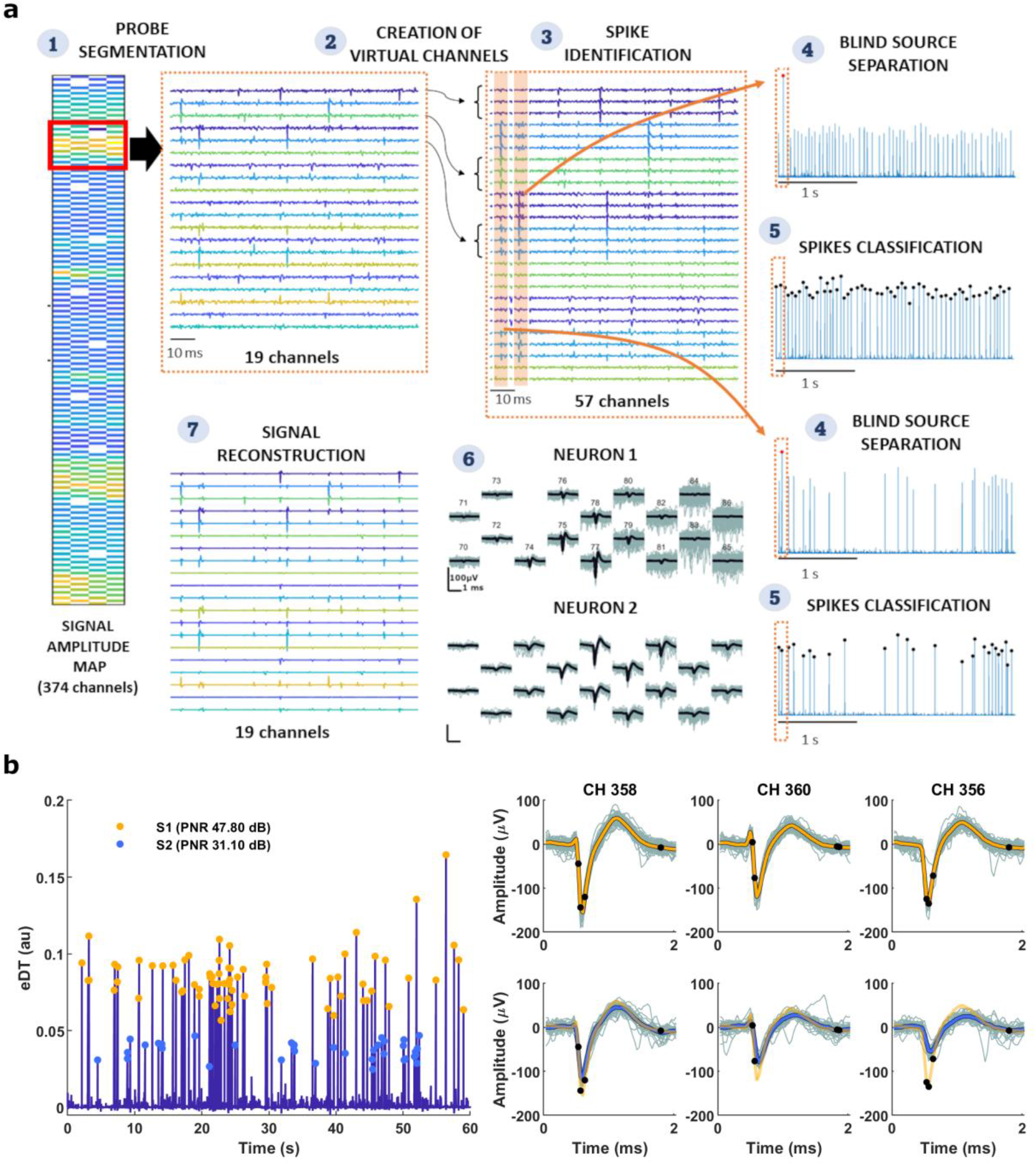
a) Main steps of the decomposition algorithm. Step 1, the dense probe is segmented into subgroups according to the signal energy (root-mean-squared) and location. The amplitude map shows the energy of each channel (color coded), the probe’s geometry, and the segmentation (red polygons). The panel on the right shows the recorded signals for the selected channels. Each line represents one channel. Step 2, virtual channels are created as delayed versions of the original data. The extension factor is 3 for clarity and graphical purposes but was usually > 6. Step 3, a peak detection algorithm locates a spike instant and initializes a separation vector for the fixed-point algorithm. Step 4, the blind source separation algorithm computes the estimated discharge train by a simple multiplication of the separation vector and the extended observations and enhances the peaks by powering to three. Step 5, the algorithm converges to the estimated source in a few iterations from which the discharge instants are recovered by a peak detection algorithm. Steps 3 to 5 are repeated N times chosen by the user. For each iteration, the algorithm can converge to a new source, an already identified source, or a non-physiological source (noise). The non-physiological and repeated sources are discarded. Step 6, the signal waveforms of each source are estimated by a spike-triggered average and the activity of all sources is summed to give the reconstructed signal (Step 7). Steps 2 to 7 are repeated for the residual signal obtained after extraction of the reconstructed signal from the original observations (peel-off procedure). The entire process is performed at each segment of the probe. b) Representative process for peak classification in a noisy estimated discharge train (eDT). The classification algorithm distinguishes two sources by analysing each detected peak in the eDT alongside the corresponding amplitudes in representative extracellular signals. Shimmer plots display the action potentials of source 1 (first row) and source 2 (second row) for the three channels. The averaged action potential of each source is showed in distinct colors (yellow for S1 and blue for S2). Black markers indicate the time points utilized in the classification algorithm.

For each spatial segment, virtual channels are created as temporally-delayed versions of each observation (step 2 in Fig. 1a). The original sources (i.e., neuronal activities) are estimated by first whitening the extended observations and then rotating them by projecting the whitened observation into a separation vector. The aim of the algorithm is to blindly estimate the separation vector of each source. For each new source to be estimated, a separation vector is initialized on the location of a detected spike (with peak detection, step 3 in Fig. 1a). The separation vector is updated iteratively with a fixed-point algorithm until convergence. The central limit theorem states that summation of sources is always more Gaussian (an indirectly less sparse) than the individual sources, thus, the criterion for the fixed-point algorithm is to maximize the temporal sparsity of the estimated sources, measured by the kurtosis (*G* = *s*^3^/3, where *s* is the estimated discharge train and G the cost function). This criterion suffices to separate the sources due to the intrinsic structure of the firing trains (most samples are zero) and the high sampling frequency of the extracellular recording. The estimated source is retrieved with a one-step matrix multiplication of the separation vector and the extended matrix of observation. The process is insensitive to overlapping action potentials, except for the case of action potentials with identical spatio-temporal waveforms, which are indistinguishable. To facilitate the detection of the peaks, the estimated source is raised to the power of three, yielding to the so-called estimated discharge train. A classification algorithm then evaluates each peak detected in the estimated discharge train with the respective amplitudes in representative extracellular signals to improve accuracy on the selection of the discharge instants (Fig. 1b). A pulse-to-noise ratio (PNR)^19^ is calculated on the estimated discharge train (see Methodology), and if the index crosses a defined threshold (25 dB), the source is confirmed, otherwise it is discarded. The PNR measures the quality of the decomposition and is an indicator of accuracy of the discharge instants identification. After identification of some sources, a peel-off procedure is applied to derive a residual signal to which the same algorithm is applied (see Methods for details).

### Performance on ground truth data

We ran the algorithm on an in vitro dataset of simultaneous loose patch and dense extra-cellular recordings (252 electrodes) from Ganglion cells in mice retina available on an online database ^20,21^. The ground-truth time instants identified from the patch signals were classified by peak detection above a manually selected threshold provided in the database.

We decomposed the initial 60 s of 14 recordings and extended the decomposition to the entire length of the recordings (see section “Extending the Decomposition” for details). The total length ranged from 180 s to 300 s. The decomposition result for a representative recording is presented in Fig. 2. The discharge trains estimated with NeuroNella from the dense extracellular recording closely followed the patch signals. To quantify the performance of the algorithm, we estimated the error rate with respect to the ground truth discharges. The error rate was computed by counting the number of false-positive and -negative discharges and dividing it by the total number of discharges from both sources. A representative false positive is shown around time 76 s and a false negative around time 102 s (Fig. 2a). Figure 2b shows the shimmer plots of the extracellular signal closest to the source triggered by the ground-truth and the discharge instants identified with NeuroNella.

**Fig. 2.**
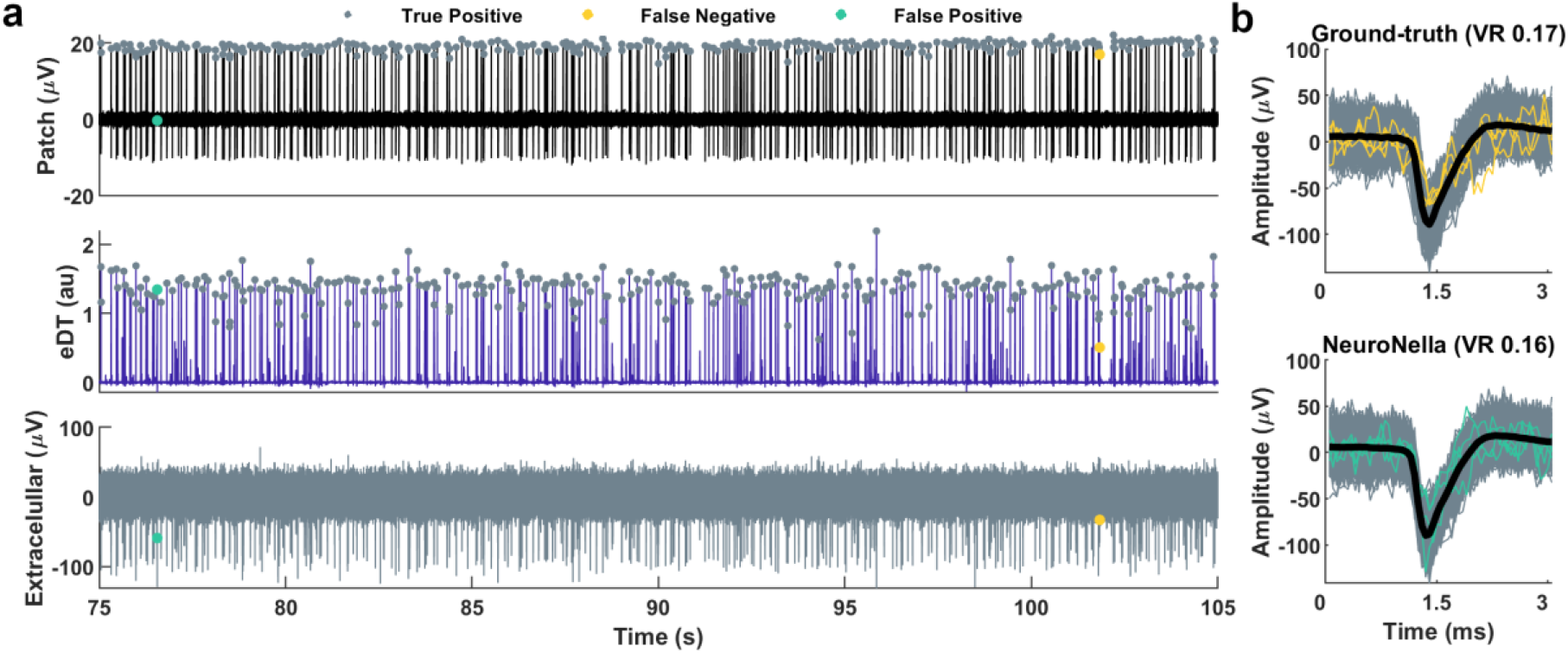
Representative comparison of the ground-truth discharge times and NeuroNella decomposition. (a) The patch signal (in black), the estimated discharge train (eDT) decomposed from the extracellular recordings with NeuroNella (in blue) and the best extracellular channel (with the largest spike amplitude, in gray) are shown for one source. Only 30 s are provided. (b) Shimmer plots for the best extracellular channel triggered by the ground-truth time instants (on the top) and triggered by the time instants detected by the decomposition algorithm (on the bottom). False-negative discharges (classified in the ground-truth data but not classified with NeuroNella) are depicted in yellow and false-positive discharges (not classified in the ground-truth but classified with NeuroNella) are depicted in green. The title contains the variance ratio (VR) of the shimmer plots.

NeuroNella achieved an averaged error rate of 0.42 ± 0.51 % (Fig. 3a), which is significantly smaller than a recent Spike Sorting toolbox^11^ that showed an averaged error rate of 5% on the same dataset. Even though a negative exponential correlation (R^2^ = 0.83) was observed between the error rates and the peak-to-peak amplitude of the action potential, similar to the Spike Sorting-based algorithms^11^, NeuroNella showcased superior performance across the entire amplitude range. This indicates that NeuroNella could automatically identify sources as small as 50 µV with an impressively low error rate, provided these sources exceeded the noise amplitude and had a unique spatial distribution. The PNR exceeded 30 dB for all estimated discharge trains (Fig. 3b).

**Fig. 3.**
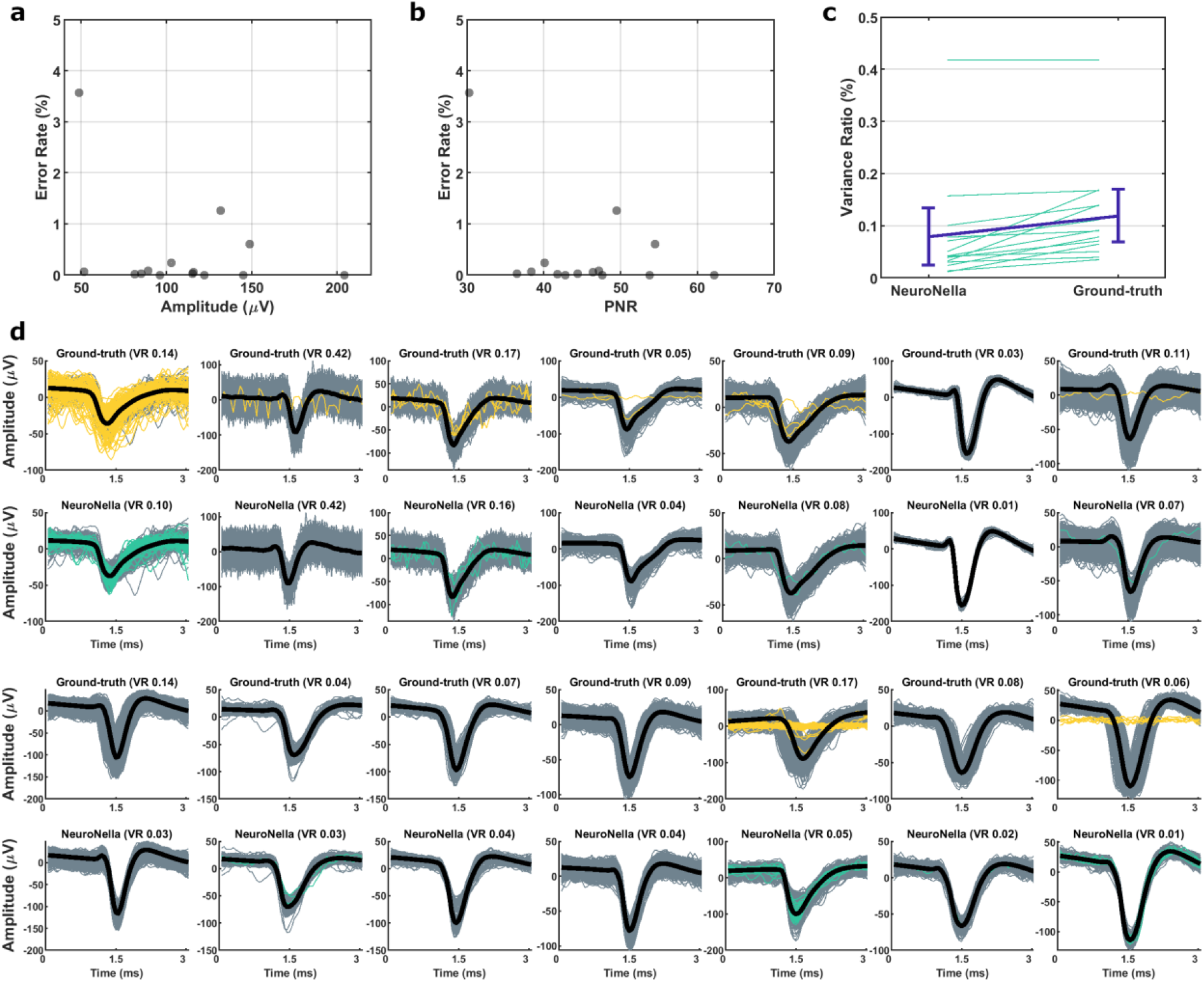
Error rate of the algorithm as a function of (a) the peak-to-peak amplitude of the decomposed source and (b) pulse-to-noise ratio (PNR) of the estimated discharge trains. Each circle represents one recording (N=14). (c) Variance ratio of the spike waveforms triggered by the original ground-truth and the discharge instants identified with NeuroNella at the extracellular recording closest to the sources. d) Shimmer plots of the extracellular recordings triggered by the ground-truth (first and third rows) and NeuroNella discharge instants (second and fourth rows) for the 14 recordings. False negatives and false positives instants are depicted in yellow and green, respectively. True positives are depicted in gray, while the averaged waveform is in shown in black. The variance ratio (VR) of each shimmer plot is presented in the title.

In order to quantify accuracy and precision in the identification of discharge instants, we computed the variance ratio of the action potential waveform (see Methods). This ratio indicates potential source mixing or deviations in the discharge instants in relate to the action potential waveform. The variance ratio was estimated for the spike waveforms triggered by the ground-truth and NeuroNella discharge instants at the extracellular recording closest to the source (largest spike amplitude). Comparatively, NeuroNella exhibited a slightly smaller variance ratio than the ground-truth (Fig. 3c), suggesting a precise selection of the discharge instants. The shimmer plots in Figure 3c demonstrate the variances within the spike waveforms. It is important to acknowledge that the ground-truth instants were derived from the patch-clamp signal, leading to slight misalignments during the triggering of the extracellular recordings, thus resulting in a slightly greater variance ratio. Additionally, Figure 3c showcases instances of false positive and false negative spikes. Interestingly, some false negative spikes (in green) closely resemble the averaged waveform (in black), while certain false positives spikes (in yellow) appear flat. These observations suggest potential misclassifications within the ground-truth instants. Specifically, in two recordings, the patch signal experienced drift and exhibited a dip in amplitude, resulting in errors during the classification of the discharge instants.

### Performance on Neuropixel data

We also tested NeuroNella on an online database that involved the use of a Neuropixel probe featuring 374 electrodes (Fig. 4a). This probe was inserted into the brain of an awake mouse spanning the posterior region and recording neural activity from the visual cortex, hippocampus, and selected thalamic areas. To facilitate decomposition, we segmented the probe into 19 subgroups, each consisting of approximately 20 channels (Fig. 4b). In total, we identified 292 neurons using NeuroNella, a lower number compared to the 492 sources provided in the dataset, which were identified by a Spike Sorting algorithm. Figure 5b reveals that the most significant disparities in source numbers were observed within segments displaying high energy. Nevertheless, the residual energy, measured as the root-mean-square difference between the original signal and the signal reconstructed from the identified neuron, exhibited only relatively minor differences between the two methods (Fig. 4c). In fact, the residual energy approached the noise level (< 7 μV, as indicated in the database) for most electrodes. Notably, the total number of discharges identified by NeuroNella algorithm exceeded that reported by the database, with 239,081 spikes against 232,071. This finding suggests that the disparity in the number of sources compared to the dataset might be attributed to the possibility that NeuroNella may have merged certain sources or that the dataset segmented some sources differently. To illustrate this further, Fig. 5 presents a high-quality source (PNR = 62.7 dB) identified by NeuroNella. This source had an 84% Rate-of-Agreement (RoA) with one source from the dataset and 5% RoA with a second source. The close resemblance of the spike waveforms identified solely by NeuroNella to the averaged waveform (Fig. 5c) strongly suggests potential misclassification of these discharges within the dataset.

**Fig. 4.**
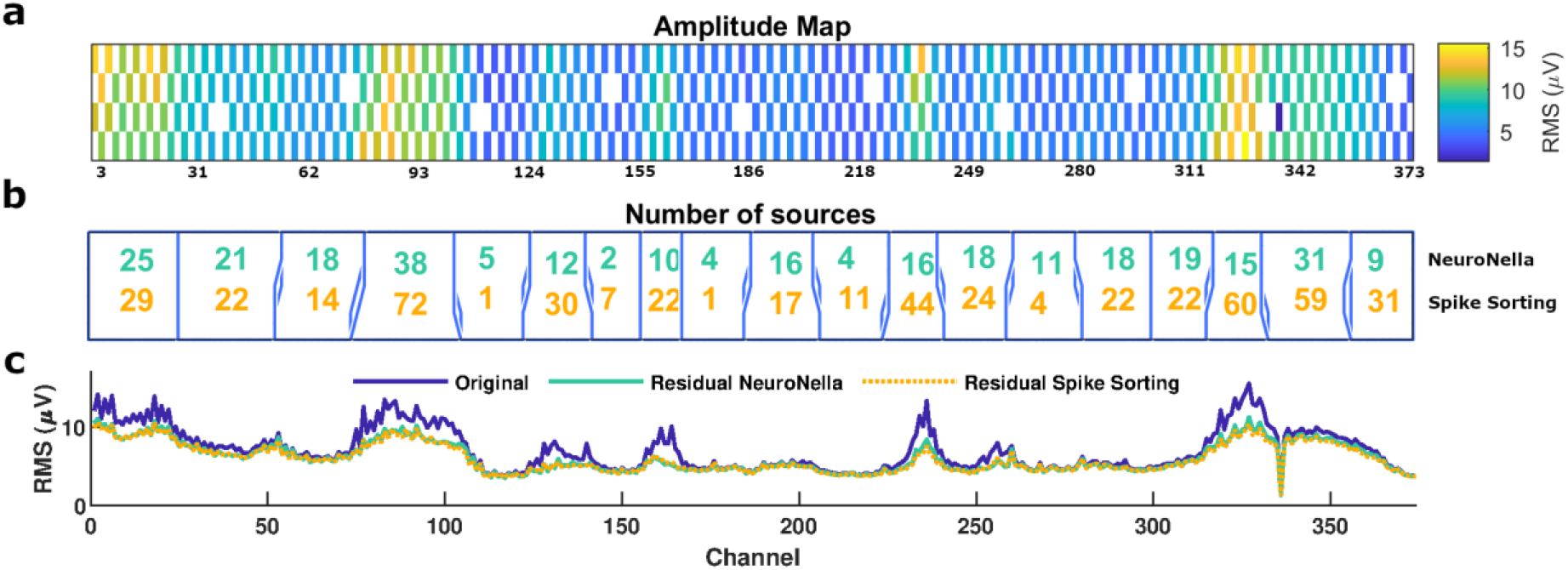
a) Amplitude map with the signal energy (root-mean-squared – colored-coded) for a 60-s duration dataset recorded with a Neuropixel probe (374 electrodes). b) Number of sources identified with NeuroNella (green) and spike sorting algorithm (orange) for each sub-group (blue polygons). c) Residual energy for NeuroNella (green) and spike sorting (orange) algorithms compared to the original signal energy (blue) for each electrode.

**Fig. 5.**
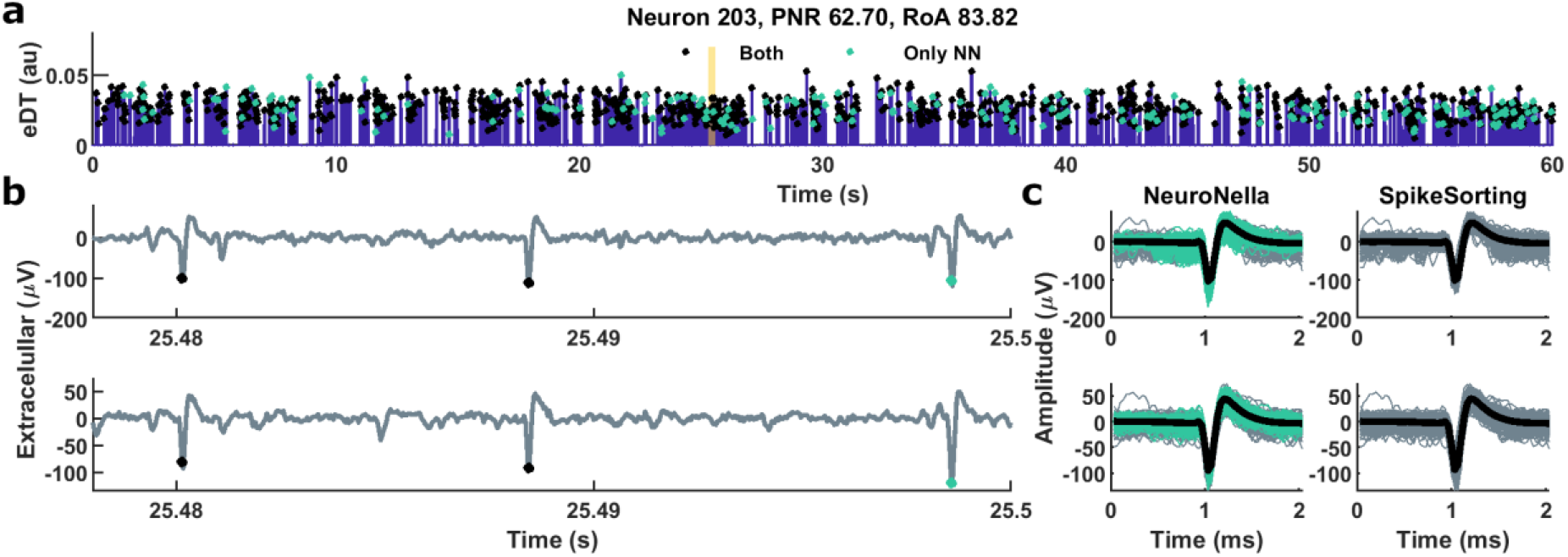
Representative comparison of the NeuroNella decomposition and the provided matched source identified by a Spike Sorting-based algorithm. a) The estimated discharge train (eDT) decomposed with NeuroNella (in blue) for the initial 60 s of recording. Discharge instants detected by both methods are indicated by black markers, while those exclusively identified by NeuroNella are marked in green. b) Zoomed-in views of the two largest-amplitude extracellular recordings aligned with the orange line in the eDT. c) Shimmer plots for the corresponding extracellular channels triggered by NeuroNella (on the left) and by the Spike Sorting time instants (on the right). Spikes classified solely by NeuroNella are depicted in green. The figure title contains the source number, the pulse-to-noise ratio (PNR) of the eDT and the Rate-of-Agreement (RoA) of the classification methods.

A total of 184 NeuroNella sources were matched to the sources provided in the database. The dataset also categorized the sources as either “good single unit” or “multiple unit”. NeuroNella identified 144 out of 223 (65%) “good single unit”, and 40 out of 269 (15%) “multiple units”. The peak-to-peak amplitude of the “single units” matched sources ranged from 18 μV to 280 μV (Fig. 6a). The sources recognized by only one method are also shown in Fig. 6a.

**Fig. 6.**
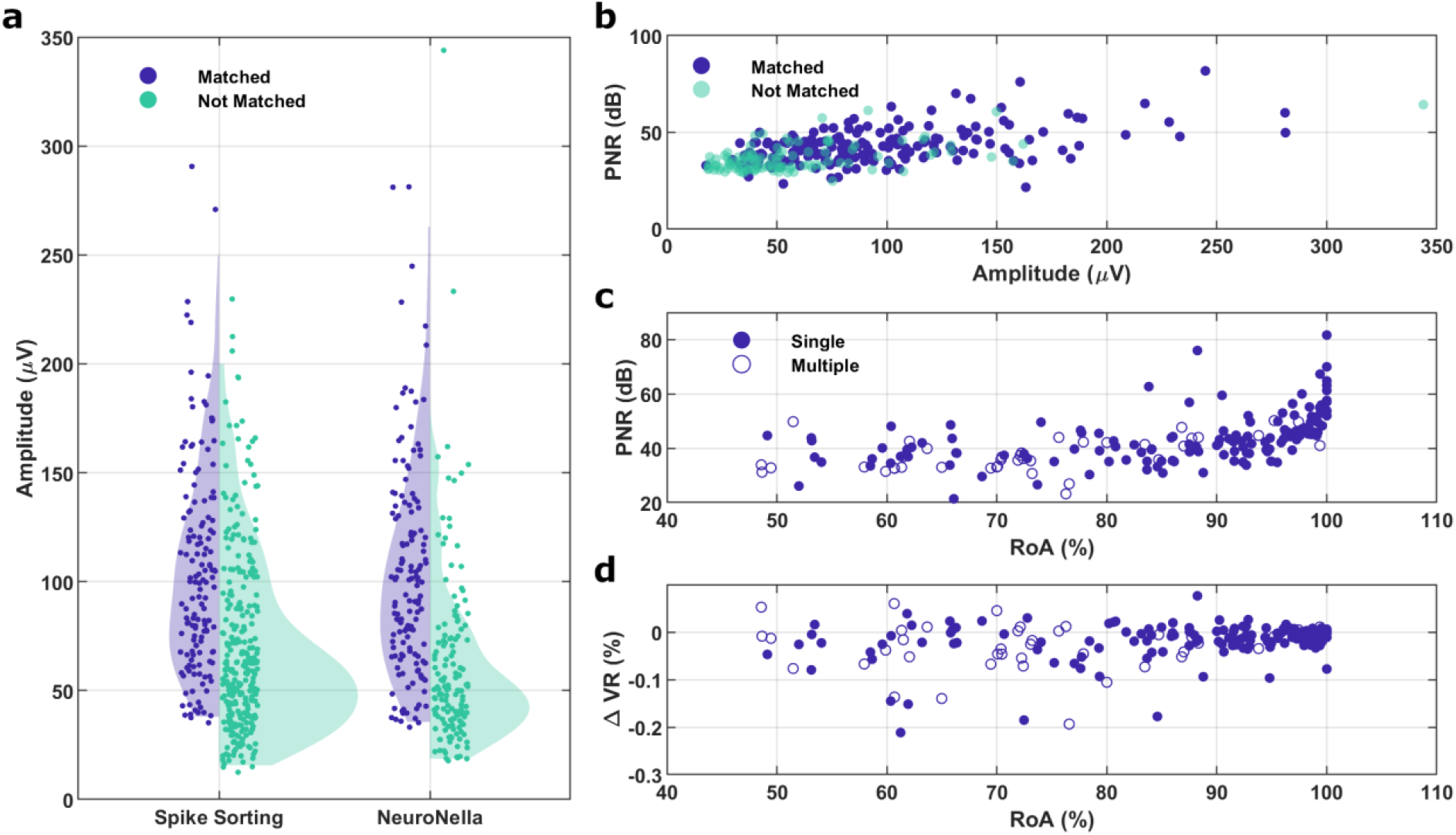
a) Peak-to-peak amplitude of the population identified with spike sorting and NeuroNella methods. Sources found by both methods (matched) are in blue and sources found by only one method are in green. b) Pulse-to-noise ratio (PNR) of the estimated discharge train in relation to the peak-to-peak amplitude of the spike waveform. c) PNR in relation to the Rate of agreement (RoA) of the spike instants for the matched sources classified as “Single Unit” (filled circles) or “Multiple Units” (open circles). d) Difference between the variance ratio of the spike waveforms triggered by the discharge instants identified by NeuroNella and Spike Sorting algorithms in relation to the RoA.

There was a correlation between the metric of quality of the blind source separation algorithm (PNR) and the amplitude of the source (Fig. 6b). Notably, most of the sources that were not matched to the dataset exhibited small amplitudes, which can also be seen in Fig. 6a. For the sources that matched, the RoA varied significantly when the decomposition quality was below 50 dB; in contrast, the RoA was 97.25 ± 1.49% (mean ± confidence interval) for sources with PNR above 50 dB (Fig. 6c). Fig. 6d shows the difference between the variance ratio of the matched sources for the spike’s waveforms triggered by NeuroNella and Spike Sorting instants. The variance ratio was observed to be smaller for NeuroNella than for Spike Sorting (median variance ratio of 0.19 vs 0.21, respectively), signifying a more precise classification.

### Collected dataset in the reticular formation

We ran NeuroNella on our own recordings from the ponto-medullary reticular formation in an anaesthetised macaque, using a 32-contact electrode (poly3, Neuronexus Inc, Ann Arbor, MI, USA). We found 26 sources in a 60 s-duration recording (Fig. 7a). The energy for the residual signals reached the noise level, indicating that the most significant sources were identified (Fig. 7b). The variance ratio was below 0.5% for all sources and was negatively correlated with the PNR (p = 0.014, R^2^ = 0.22, Fig. 7c).

**Fig. 7.**
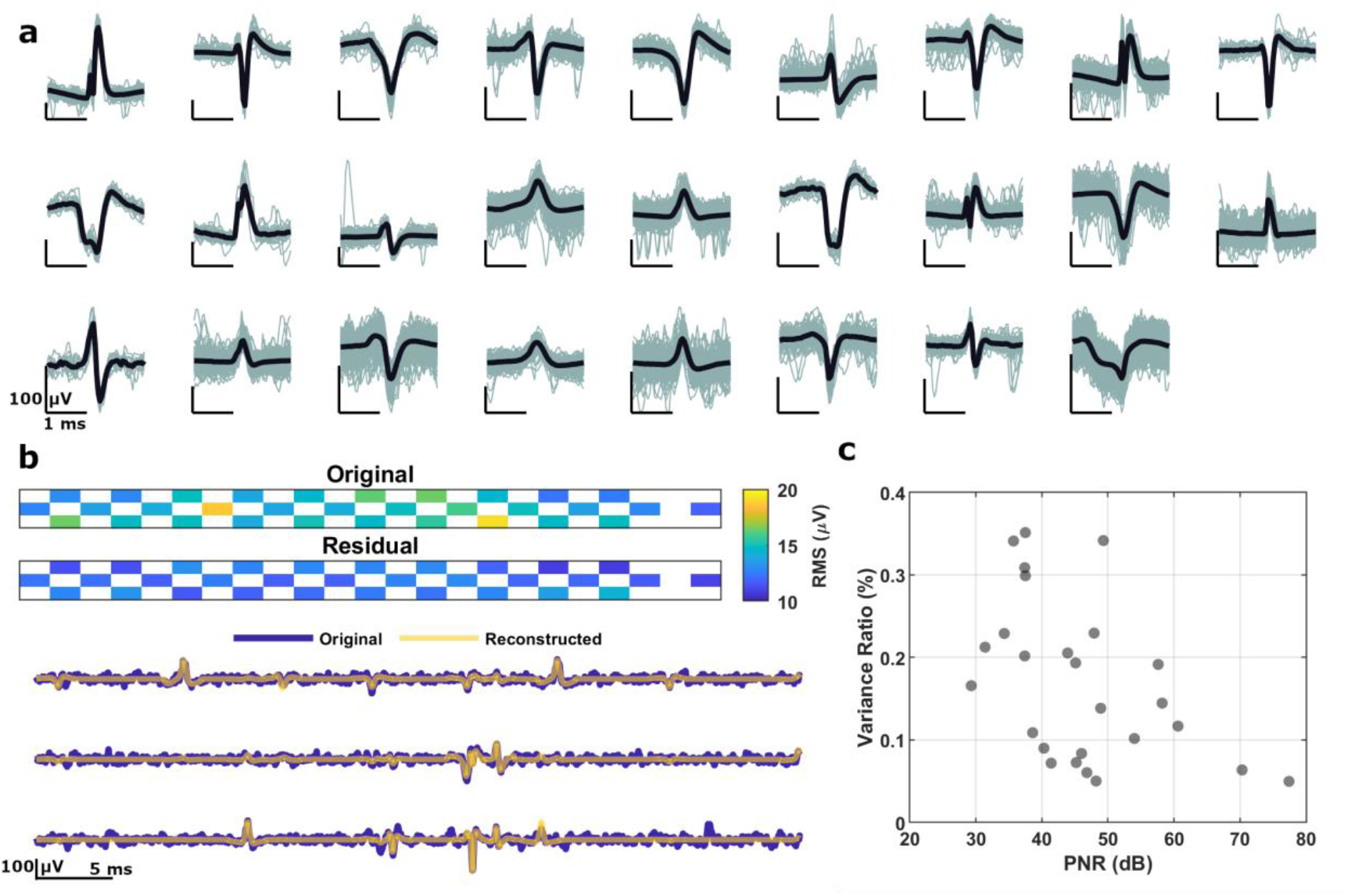
a) Shimmer plot of 26 sources identified with NeuroNella in an extracellular recording of the reticular formation. b) Amplitude map with the signal energy (root-mean-squared – colored-coded) for a 60-s duration dataset recorded with a linear U probe with 32--electrodes, and the residual energy for comparison. Bellow, representative signals recorded in three electrodes (blue) and the respective reconstructed signals from the identified neurons (yellow). Bellow, the c) Variance ratio as a function of the pulse-to-noise ratio (PNR) of the estimated discharge train.

We then extended the decomposition to 20 min and found 10 neurons constantly active. Notably, the discharge identification for the extended decomposition was robust to recording instability, as shown by drifts in the amplitude of the extracellular signals (Fig. 8a) and changes in the action potential waveforms with time (Fig. 8b).

**Fig. 8.**
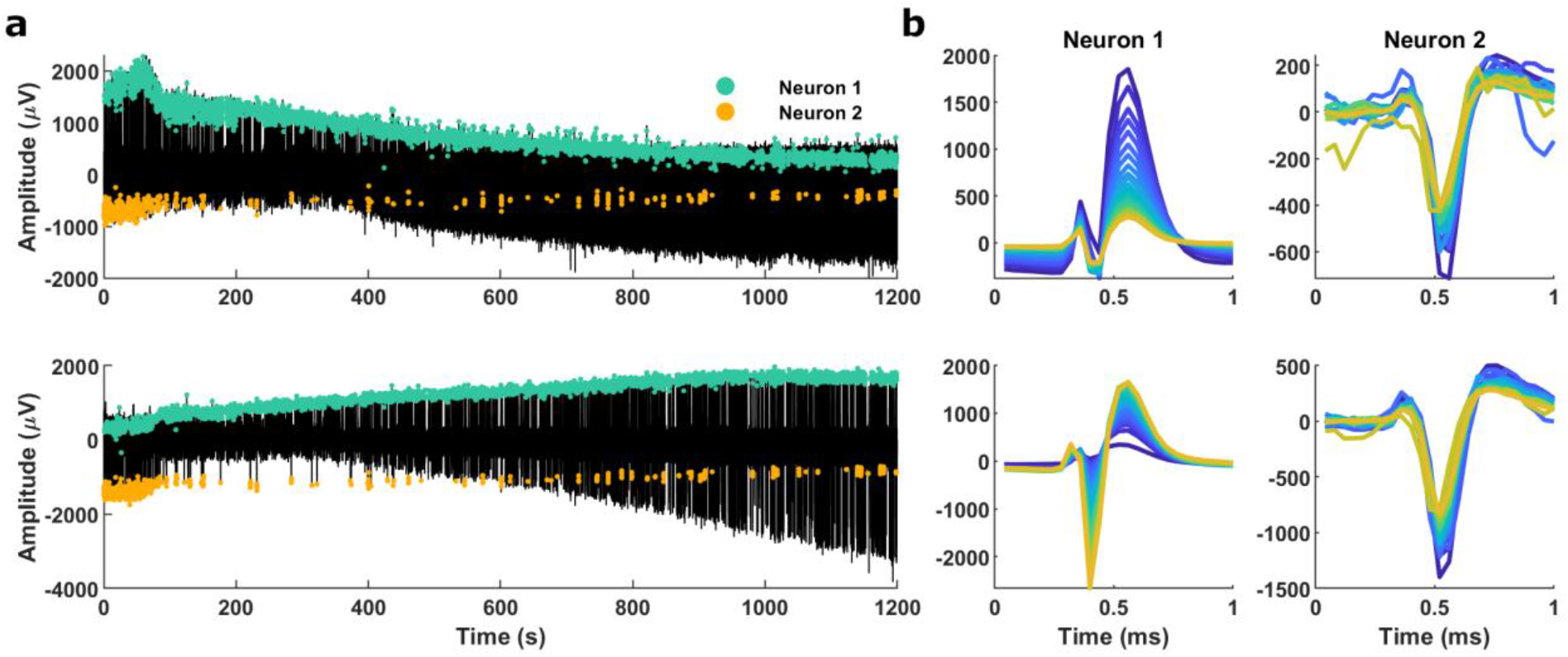
Representative results for the decomposition of a 20-min recording. a) Two extracellular signals displaying the discharge instances identified for two neurons (green and orange markers). b) Action potential waveforms corresponding to Neuron 1 (on the left) and Neuron 2 (on the right) for the respective extracellular signals estimated at every minute of recording.

## DISCUSSION

We developed a method that leverages the sparsity of the action potentials from multichannel electrode arrays implanted in different locations in the nervous system of mice, rats, and monkeys. NeuroNella shows high reliability in identifying the same spike even in cases of overlapping action potentials and shows a consistent identification of the same neuron. We found matching accuracy between the patch-clamp and multichannel extracellular action potentials close to 100%. The spike times of the neurons were identified in a fully automatic way, therefore, opening new avenues not only for offline processing in basic and applied neuroscience research but also for future online use in brain/machine interfaces.

The automated decomposition and extension processes demonstrate remarkable efficacy in detecting sources of varying amplitudes, exhibiting an exceptional level of accuracy. We observed that error rates > 0.5% were predominantly associated with the challenges of peak identification within the estimated discharge train and the patch-clamp signal. In Figure 3, numerous supposedly false negative discharges do not resemble the action potential waveform, while the supposedly false positives did, possibly underscoring the impact of utilizing a simplistic amplitude threshold for identifying discharge instants within the patch-clamp signal. Thus, our findings also highlight the significance of reliable ground-truth data, serving as a critical reference point for error validation and algorithm refinement^21^.

Through our analysis, we found that adopting a more rigorous approach favoring decompositions with a PNR exceeding 50 dB guarantees the overall reliability of our results. The PNR index, as defined in this study by Equation 10, derives from the PNR index of the decomposition of high-density electromyogram into motor units spike trains^19^. For the later application, the threshold is usually set to 30 dB, as it provides sensitivity higher than 90% and errors below 2%^19^. In this study, the PNR values and the recommended threshold are notably higher due to a distinct calculation method – employing a power of three instead of the power of two used in the electromyogram, which further nonlinearly amplifies the peaks over the noise. In instances where the estimated discharge trains exhibit a low PNR, typically associated with sources characterized by small amplitudes, close proximity to other active sources, or limitations in the spatial resolution of the acquisition system, we strongly recommend a manual inspection with the semi-automatic classification algorithm (Fig. 1b). This algorithm is specially designed to consider the distinctive waveform characteristics of each spike across neighbouring electrodes to classify the peaks identified in the estimated discharge train properly.

A critical step in the NeuroNella decomposition involve segmenting the probe into smaller subgroups of electrodes. However, this segmentation process did not consider the spatial distribution of individual sources; instead, it relied on overall signal energy, potentially resulting in suboptimal segmentation regarding the selectivity of individual sources. Despite this, the decomposition process proved effective, highlighting its resilience even with incomplete spatial information. A follow-on study may evaluate if probe segmentation based on detected neuronal activities increases the number of identified sources and improve the overall quality of the decomposition.

Results indicate that the comparatively lower number of sources identified by NeuroNella appears to be more closely associated with the intricacies of spike identification rather than a fundamental limitation of the Blind Source Separation algorithm in detecting some sources. Several key observations support this hypothesis. In Figure 6, NeuroNella successfully detected sources within the same amplitude range as the spike sorting provided with the online dataset. Additionally, Figure 5 illustrates possible misclassifications by that Spike Sorting algorithm, suggesting subdivision of sources in the dataset. Finally, Figure 6d exhibited a lower variance ratio of the spikes when triggered by the NeuroNella discharge times compared with the times provided in the dataset. Notably, considering a minimal 0.03% difference between the variance ratio of matched sources, NeuroNella displayed lower values for 90% of the sources. This suggests that it is more likely that Spike Sorting missed spikes (or split sources) rather than that NeuroNella falsely selected pulses (or merged sources), since otherwise the variance ratio would have been higher. These misclassification cases became particularly evident in decompositions characterized by a high PNR and a low RoA. Therefore, sources featuring PNR values above 40 dB and with RoAs ranging between 60% and 90% in Figure 6b likely resulted from erroneous classifications by the Spike Sorting algorithm.

In addition to the potential misclassifications, it is important to highlight that the unmatched sources in the online dataset were predominantly classified as “multiple units”, which might not hold significant relevance for many research studies. NeuroNella seemed to prioritize the automatic identification of “good single units”, which will ensure a more focused and relevant dataset for further analysis.

Our investigation further demonstrated that the NeuroNella algorithm effectively extended the decomposition process to large datasets (30 min). Results showed high robustness of the extended decomposition process in handling minor probe shifts (Fig. 8). This resilience underscores the potential application in longitudinal studies and prolonged recordings sessions. Moreover, the algorithm holds promise for online decomposition as it is based on a simple multiplication of the separation vector (identified in an offline step) and the new data. The process follows the same principle of online decomposition of high-density electromyogram ^22,23^. However, it is important to note that this process only guarantees precision for the high-energy sources, and not for smaller sources identified after the peel-off procedure.

## METHODS

### Details of the Algorithm

In this section, we describe a general framework of a blind source separation algorithm for extracellular recordings. It starts by describing the segmentation procedure to optimize the decomposition of dense probes; it then proceeds to the data representation of electrophysiological recording as an instantaneous convolutive mixture model ^17,18,24–26^; and finishes with the theoretical basis for the separation algorithm based on a combined approach of independent component analysis^27^, fast independent component analysis (fastICA,^24,28,29^), convolutive blind source separation algorithm ^17^ and peel-off procedure ^30^.

### Segmentation of dense probes

The neuronal activity can be considered spatially sparse in a dense probe, as usually few electrodes register the electrical signal, depending on the electrode density and its proximity to active sources. Observations that do not register activity of the same source unnecessarily increase the computational cost and may lead to the convergence of the Blind Source Separation algorithm to other more significant sources (see Data Model section). Segmentation into sub-groups of channels is therefore crucial for large probes. In cases of more than 40 channels, we segment the probe into sub-groups of neighbouring electrodes with similar signal energy (root-mean-square). For this, a K-means++ algorithm uses the energy recorded in each electrode and its location in the probe (X and Y) to segment it into N non-overlapping sub-groups. The number of segments (N classes) depends on the electrode density in the probe, but generally aims at having about 25 channels per sub-groups, as it offers a good trade-off between the total processing time and the number of identified sources. It should be noted that the source can still be identified even when its spatial distribution is divided into two or more sub-groups, provided it has a unique spatial-temporal distribution and a significant signal energy compared to other sources. Nonetheless, decomposition of overlapped segments may alleviate potential subdivision effects. In both cases, duplicity in the source identification may occur, but is further eliminated.

### Data Model and Separation Background

The problem we want to address is the blind identification of neuronal activities extracted from electrophysiological recordings. It models like a classical cocktail-party problem, where the intent is to retrieve the speech signals of multiple speakers who are simultaneously talking in a room with multiple microphones. The speech signal is filtered by the acoustic properties of the environment before reaching each microphone, yielding a unique spatial distribution of each speech. Likewise, the electrophysiological signals recorded by implanted electrodes in brain tissue are the filtered versions of the action potential trains of multiple neurons.

The electrophysiological data can be modelled as an instantaneous convolutive mixture, as it consists of the convolutional combination of action potential waveform and the discharge times. Assuming the source signals arrive at the sensors at the same time, the model is also said to be instantaneous. The same model is found in the case of invasive and non-invasive high-density electromyogram extensively described elsewhere ^17,18,24^. The observations (channels) comprise the summation of action potential trains in whose waveform is considered invariant in the time and specific for the neuron and channel. Assuming N active neurons are recorded by M electrodes, the signal on each channel can be described as:

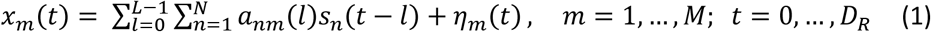

Where *x*_*m*_(*t*) is the observation in the *m*^*th*^ channel at the time instant *t* (in the discrete time), *D*_*R*_ is the duration of the recording (in samples), *a*_*nm*_(*l*) is the action potential waveform of the *n*^*th*^ neuron in the *m*^*th*^ channel, *L* is the length of the action potential waveform, *s*_*n*_ is the discharge train of the *n*^*th*^ neuron modelled by a series of Dirac Delta function (*s*_*n*_(*t*) = ∑_*k*_ *δ*(*t* − *T*_*n*_(*k*)), where *T*_*n*_ are the discharge instants), and *η*_*m*_(*t*) is the additive noise at channel *m*. Due to the refractory period, we can assume no overlapping of action potentials for the same neuron, thus *T*_*n*_(*k* + 1) − *T*_*n*_(*k*) > *L* for *∀k ∈* ℕ.

In the vector-matrix notation, Equation (1) becomes

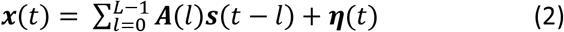

Equation (2) is a finite impulse response (FIR) model, where ***x***(*t*) = [*x*_1_(*t*), *x*_2_(*t*), … , *x*_*M*_(*t*)]^*T*^ is the vector of observations, ***A***(*l*) = [***a***_**1**_(*l*), … , ***a***_***N***_(*l*)] is the action potential filter of order L and size *M x N*, and ***s***(*t*) = [*s*_1_(*t*), … , *s*_*N*_(*t*)]^*T*^ is the vector of sources at time instant *t*. Superscript ^*T*^ denotes transposition of a vector or a matrix.

The simplest approach to separate a convolutive instantaneous mixture is to represent it as a standard linear model by its compact form. First, the convolutive model is rewritten in its matrix form

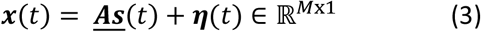

Where ***<u>A</u>*** = [***A***(0), ***A***(1), … , ***A***(*L* − 1)] *∈* ℝ^*M*x*NL*^, and ***<u>s</u>***(***t***) = [***s***^*T*^(*t*), ***s***^*T*^(*t* − 1), … , ***s***^*T*^(*t* − *L* + 1)]^*T*^ *∈* ℝ^*NL*x1^ contains the latest L samples of the sources.

By using a sliding window of length K+1, here after referred to as the extension factor, it yields to a block-based model

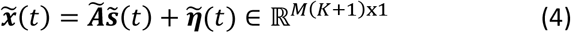

Where, 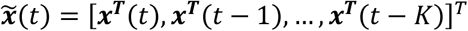, is the extended vector of observations containing the original signals and K delayed versions, and

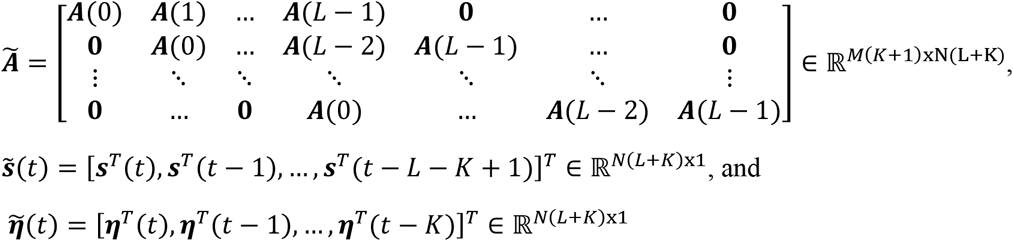

The mixing matrix ***Ã*** is a generalized Sylvester block Toeplitz matrix, having *K* + 1 vertical and *L* + *K* horizontal *M*x*N* blocks, and is full column rank ^31^.

Conversely to spike sorting algorithms, the blind source separation algorithm does not retrieve the action potential waveforms *per se*. The independent sources are blindly estimated based on the spatiotemporal statistics of the mixing matrix and the discharge trains (detailed shortly in this section). This is accomplished by finding a corresponding demixing system of Equation 4, as given by Equation 5.

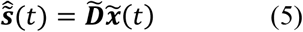

Where 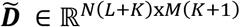 is the matrix of coefficients of the separating system filters, also called matrix of weights, and 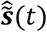 is the estimated source vector. The demixing problem builds on the fundamental assumption of statistical independence or sparseness (non-Gaussianity) of the sources. The model also assumes that the source signal vector are zero-mean stationary real signals and is independent of the noise vector. To make the zero-mean assumption hold we center the observation matrix by subtracting their sample mean. Moreover, the extension factor (K+1) should be large enough to assure *M*(*K* + 1) ≥ *N*(*L* + *K*) to yield to an overdetermined multiple-input multiple-output demixing model. One must account for the dimension growth of the problem when the extension factor and/or the number of channels (M) are too large, especially for recordings with high sampling frequency (20 kHz) and long duration, as it leads to prohibitively high dimensions and computation overload. An extension factor (*K* + 1) = 300/*M* was used, where M is the number of channels per sub-group of the probe.

Once the convolutive mixture model (Equation 1) is approximated to an instantaneous linear mixture model (Equation 5), we can proceed to the separation problem by standard linear independent component analysis. The approach chosen was fast independent component analysis (fastICA) ^32^, as it has been successfully applied and validated for high-density electromyogram ^24^.

For mathematical convenience, the zero-mean extended observations are spatially whitened, such that the channels are mutually uncorrelated and have unit variance. To ensure stability, a regularization factor equal to the 20^th^ percentile of the eigenvalues was used. After the transformation, and omitting the noise for simplicity, it gives

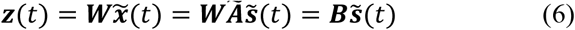

Where ***W*** is the whitening matrix, and ***B*** is an orthogonal matrix owing to the assumption of independence of sources. The demixing model is then given by

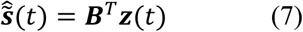

Equation 7 holds that source separation is achieved based on the estimation of a separation matrix ***B***. This matrix is found by an iterative procedure (fastICA) that optimizes a contrast function chosen based on the statistical properties of the neuronal discharge trains: its intrinsically sparse distribution (most samples are zeros), closely related to supergaussianity. According to the central limit theorem, the sum of non-gaussian random variables is closer to gaussian than the original ones. Therefore, each source of Equation 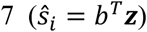 can be iteratively estimated by finding the vector *b* that maximizes the nongaussianity of *b*^*T*^***z***. Estimates representative for the statistical properties of the source and related to nongaussianity can be used as contrast functions, such as kurtosis and skewness given by *G*(*s*) = *s*^4^/4, and *G*(*s*) = *s*^3^/3, respectively ^33,34^. The separation vector is optimized iteratively by a fixed-point algorithm given by ^32^:

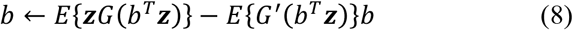

Where *E* stands for mathematical expectation. Normalization of the separation vector to unit norm is necessary at each iteration to keep the variance of the estimated source constant. This loop runs until there is no significant changes in the separation coefficients, such that 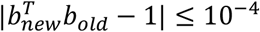, or up to 30 iterations.

To facilitate the identification of the discharge instants on the estimated source (*ŝ*_*i*_ = *b*^*T*^***z***), we enhance the prominence of the peaks by calculating the estimated discharge train, as follows.

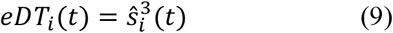

The peaks are then detected by a first-derivative method and classified by a clustering method (see Peak Classification Algorithm section). The quality of the classification of the peaks is evaluated with the modified pulse-to-noise ratio index ^19^.

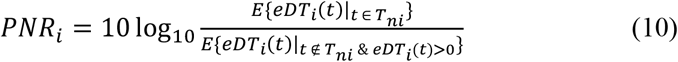

Where *E* stands for mathematical expectation and *T*_*ni*_ represents the selected discharge instants of source *i*. Note that only positive noise are considered in the denominator. PNR indicates accurate peak identification, as high PNR is achieved when the base-line noise levels are low in comparison to the amplitude of the selected peaks. The source was accepted if PNR was greater than a threshold of 25 dB.

We then compute the action potential waveforms of the source for each channel with spike-triggering average (STA) of the original recordings, and the entire action potential trains are recovered by convolving the action potential waveforms with the pulse train.

The process described above is the basis for recovering one independent source. For multiple sources, the fixed-point algorithm must be run several times. For every source identified, we check for duplicity and, if false, the separation vector, the discharge instants and the waveforms are stored. Two sources are considered equal if the cross-correlation between the discharge trains normalized by the minimum number of discharges is higher than 0.5, accounting for time shifts of the discharge instants of ±0.1 ms. When duplicity occurs, the source with the smallest PNR is deleted. To avoid the convergence of the algorithm to sources already found, a deflationary orthogonalization procedure 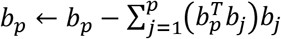 is added in the loop following the update step (Equation 8) and prior to normalization. Duplicity is also avoided by choosing adequate initialization of the separation vector. A good initialization is the observation vector at the time instant of a detected discharge, thus, *b*(0) = ***z***(*t*_*k*_), where ***z*** is the whitened extended observations and *t*_*k*_ is the detected time instant ^17^. The time instants are chosen based on an activity index, computed as the summation of the squared whitened observations ^17^, as high values indicate neuronal activity from at least one source recorded by at least one channel. Once the discharge train of a source is identified, the corresponding time instants of the activity index are zeroed, and the maximum value of the activity index sets the initialization vector for the next source.

The fastICA, as any blind source separation algorithm, is limited to extract the sources that most contribute to the signal. To allow extraction of more sources, the peel-off procedure ^35^ is performed to remove from the original data the action potential trains of the extracted sources, allowing the convergence of the fastICA to less significant sources. This procedure may impose minor estimation errors in the model if repeated, which did not appear to occur here.

In sum, the algorithm behind NeuroNella is to segment the probe into sub-groups and perform two steps of fastICA, one with the original electrophysiological recordings and the other with the residual signal following the peel-off procedure. In each step of fastICA, a set number of iterations determines how many times the algorithm will attempt to extract a source by initializing a separation vector and optimizing it with the fixed-point algorithm. Duplicity of neurons are checked at each step of the fastICA and for adjacent sub-groups.

### Peak Classification Algorithm

Some estimated discharge trains have poor signal-to-noise ratio due to similarities in the spatial distribution of nearby neurons. High background noise may lead to misclassification of the discharge times and necessitates further inspection. To overcome this issue, we developed an algorithm automatically to recognize the peaks in a noisy estimated discharge train based on a clustering analysis that accounts both for the amplitude of the peaks and the corresponding amplitude in three extracellular signals closest to the source (taken from the separation vector *b*).

The algorithm is based on a hierarchical clustering algorithm limited to 20 clusters (function *clusterdata*, from Matlab®). First, the algorithm detects peaks in the estimated discharge train with minimal distance of 1 ms, thereby selecting both signal and noise. Then, the algorithm computes the associated waveforms of each selected time instant by spike-triggering three extracellular signals with a window length of 3 ms. Reference waveforms are derived by averaging only the waveforms corresponding to peaks surpassing a specific threshold, typically set to 0.5% of the second norm of the estimated discharge train. As an input for the clustering analysis, at each time point, the algorithm calculates the root-mean-square deviation between the associated waveform and the reference waveforms and assesses the waveform amplitude at four critical points: the peak and three points automatically selected for their significant variability among all waveforms (Fig. 1b). In the final phase, the algorithm arranges the clusters based on the estimated discharge train amplitude’s centroid and explores reducing the number of clusters by identifying similarities in the action potential waveforms of all clusters with the first one. Clusters are merged under the following conditions: i) the combined correlation of the action potential across the three extracellular signals exceeds 2.5; ii) the centroid of the discharge train amplitude surpasses 0.5 of the first cluster’s centroid. iii) the root-mean-square deviation in the action potential for the best extracellular signal falls below 3. The cluster with the highest estimated discharge train amplitude is stored.

This algorithm is adaptable for semi-automatic use during manual inspection of peek classification. The user has the flexibility to define the reference waveforms (based on *a priori* discharge identification), the number of clusters, the number of critical time points in the waveform’s amplitude, and the weighting of each variable inputted into the hierarchical clustering. Additionally, users can set the criteria thresholds for cluster merging, facilitating adaptability and customization to specific noisy estimated discharge train.

### Extension of the Decomposition

NeuroNella can run on a computer without GPU, however, the size of the extended observation matrix is a limiting factor. The computational cost is determined by the number of selected channels, the extension factor, the time duration, and the sampling frequency. In this study, decompositions were limited to recording segments ranging from 30 s to 120 s. If a longer decomposition is necessary, the decomposition of one segment can be extended to the next segment of data. To account for minor drifts in the probe and subsequent changes in the action potential waveform and spatial distribution of each source, it is recommended to perform the extension procedure in segments no longer than 60 s of recording, with an overlap of 50%, and to update the estimation parameters based on the new data. The extension procedure used here is similar to the online decomposition of high-density EMG described in^22^. A slightly modification of Equation 7 allows the retrieval of the estimated discharge train without whitening the new data, as follows ^17^:

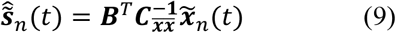

Where 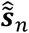 is the estimated sources for the new segment and 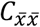 stands for the covariance matrix of the extended observations, updated at the covariance matrix of each new segment 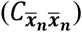 by Equation 10.

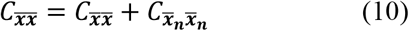

Every separation vector that constitutes the separation matrix (***B*** = [*b*_1_ … *b*_*m*_]) is obtained from the initial decomposed segment, as follows:

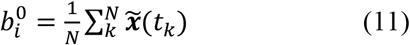

Where *t*_*k*_ is the *k’*th discharge time (from 1 to N discharges) of the *i*’th source extracted.

The process of identifying discharge times within the newly estimated discharge train involves the hierarchical clustering analysis (described in “Peak Classification Algorithm” section) of a data segment that overlaps with the preceding segment. This overlapping segment provides a reference to historical discharge times and action potential waveforms. This process compensates for potential drifts in the recording system and accommodates the activation of new sources, both of which contribute to fluctuations in the amplitude of both signal and noise in the estimated discharge train.

The newly identified discharges are used to update the separation vector, as follows:

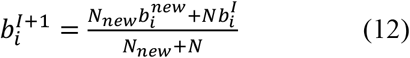

Where, *N*_*new*_ is the number of identified discharges in the new segment, 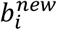 is the separation vector evaluated within the new segment, and *N* denotes the cumulative count of identified discharges up to the current segment.

Since the Peak Classification Algorithm can also identify new sources contributing to the estimated discharge train (as seen as Source 2 in Fig. 1b), those exhibiting a PNR greater than 25 dB are stored and subsequently examined in the new segment.

### Experimental Recording

One unpublished dataset was analysed, gathered in the Baker laboratory in Newcastle University. This was recorded from the medullary reticular formation in a terminally anaesthetised female macaque monkey. Under inhalation anaesthesia (1-2% sevoflurane), a craniotomy was made in the skull adjacent to the foramen magnum, and a laminectomy made of the cervical spinal cord. The head was then fixed in a stereotaxic frame, angled down at 65°; the spinal cord was clamped using spinal clamps to be approximately horizontal. Anaesthesia was then switched to an intravenous infusion of ketamine (6 mg/kg/hr), alfentanil (0.2-0.3 μg/kg/min) and midazolam (0.14 mg/kg/hr), which we have found to provide a stable anaesthetic plane whilst maintaining neuronal activity. The cisterna magna was opened to visualise the obex. Recordings were made with a silicon probe electrode (poly3, Neuronexus Inc, Ann Arbor, MI, USA) which had 32 contacts (diameter 15 μm) arranged in three columns, with interelectrode spacing 25 μm and total length of the recording array 290 μm (see ^36^Kraskov et al, 2020 for an image of the array layout). The electrode was held in a stereotaxic manipulator angled rostrally at 35° relative to the vertical line perpendicular to the spinal cord. The electrode tip was first zeroed to the obex landmark and was then moved to penetrate into the brainstem 2mm rostral, 1mm lateral to obex. Recordings were made at a penetration depth of 8.2 mm below the depth measured at obex, which should lie within the *nucleus gigantocellularis*. Signals from all 32 channels were captured using an Intan RHD2132 headstage amplifier and associated USB interface (Intan Technologies Inc, Los Angeles, CA, USA), with sampling rate of 25 kHz and bandpass 1Hz-10kHz.

## Data Processing

### Ground truth data decomposition

The online dataset provides 19 simultaneously recordings of extracellular and patch-clamp signals from ganglion cells of the retina. However, four recording showed bad patch-clamp signals (signal-to-noise ratio < 3.3 dB) which made it inappropriate to be taken as ground-truth. For 15 recordings with satisfactory signal-to-noise ratio, the decomposition process started by segmenting the 252-channel probe into sub-groups of approximately 36 channels, and by taking the sub-group closest to the patched source (best channel provided in the database). We ran the decomposition algorithm for the first 120 s and extended the observation matrix by a factor of 5 (K = 5, empirically found). Every source extracted was compared to the ground-truth spike times, and if the error rate was smaller than 20% the algorithm ceased. The peeling-off procedure was not necessary. We then extended the decomposition to the entire length of recording. A visual inspection on the ground-truth data was performed and the threshold level was manually adjusted to correct some misclassified spikes.

### Neuropixel data decomposition

We ran NeuroNella in the first 60 s of the Neuropixel dataset. The probe was segmented into 19 sub-groups with approximately 20 channels each. The extension factor was set for each subgroup as the ceiling integer that yielded an extended observation matrix of 300 virtual channels (extension factor multiplied by the number of channels). The number of iterations in each step of the decomposition for the original and peeled-off signals was set to 80, allowing us to find up to 160 sources in each segment of the probe.

### Reticular formation data decomposition

For the reticular formation recording, we decomposed the first 60 s of data for the single segment comprising 32 channels. The extension factor was 12 and the number of iterations was 80 for each step of the decomposition. The decomposition was then extended for the first 30 min of recording following the procedures described in the section “Extension of the Decomposition”.

## Data Analysis

For each extracted source, we obtained the spike waveform at all channels with spike-triggered averaging and the activation map, computed as the peak-to-peak amplitude of the spike waveforms across the entire probe. We also estimated the spike train of each source at each channel by convolving the spike waveforms with the discharge times. The residual signal was then obtained by subtracting the reconstructed spike trains from the original data. The residual energy was found as the root-mean square of the residual signal.

The accuracy of the discharge times was estimated by the variance ratio (Equation 13)^37^, which indicates the variability in the action potential shape across the spike train.

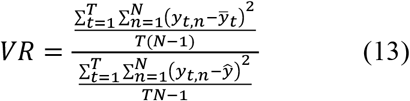

Where T is the number of time points, N is the number of discharges, *y*_*t*,*n*_ is the value of the *n*^*th*^ discharge at time point, 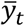 is the spike waveform value at time point *t*, averaged over the N discharges, and *ŷ* is the grand mean.

To assess the performance of NeuroNella, we estimated the Error Rate of the extracted sources in comparison to the ground-truth sources. The error rate was computed by counting the number of false-positive and -negative discharges and dividing it by the total number of discharges from both sources. A tolerance of 0.5 ms of deviation was considered acceptable in the analysis.

For the Neuropixel sources, we first matched all source identified with NeuroNella to the sources provided in the database. Sources were considered matched if the cross-correlation between spike trains was higher than 0.5. We only analysed sources with at least 10 discharges in the 60-s segment. Instead of error rates, we compared the discharge times of the NeuroNella and Spike Sorting matched sources by the Rate of Agreement, as follows:

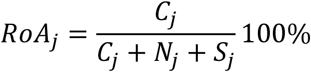

where *C*_*j*_ denotes the number of discharges of the *j*th source identified by both methods, and [*N*_*j*_, *S*_*j*_] are the number of discharges identified only by NeuroNella or Spike Sorted methods, respectively. The discharge time tolerance was 0.5 ms.

