## Supplementary figures and images for "NeuroNella: A Robust Unsupervised Algorithm for Identification of Neural Activity from Multielectrode Arrays"

### Supplemental Figures

SUPPLEMENTARY MATERIAL

Figure A1

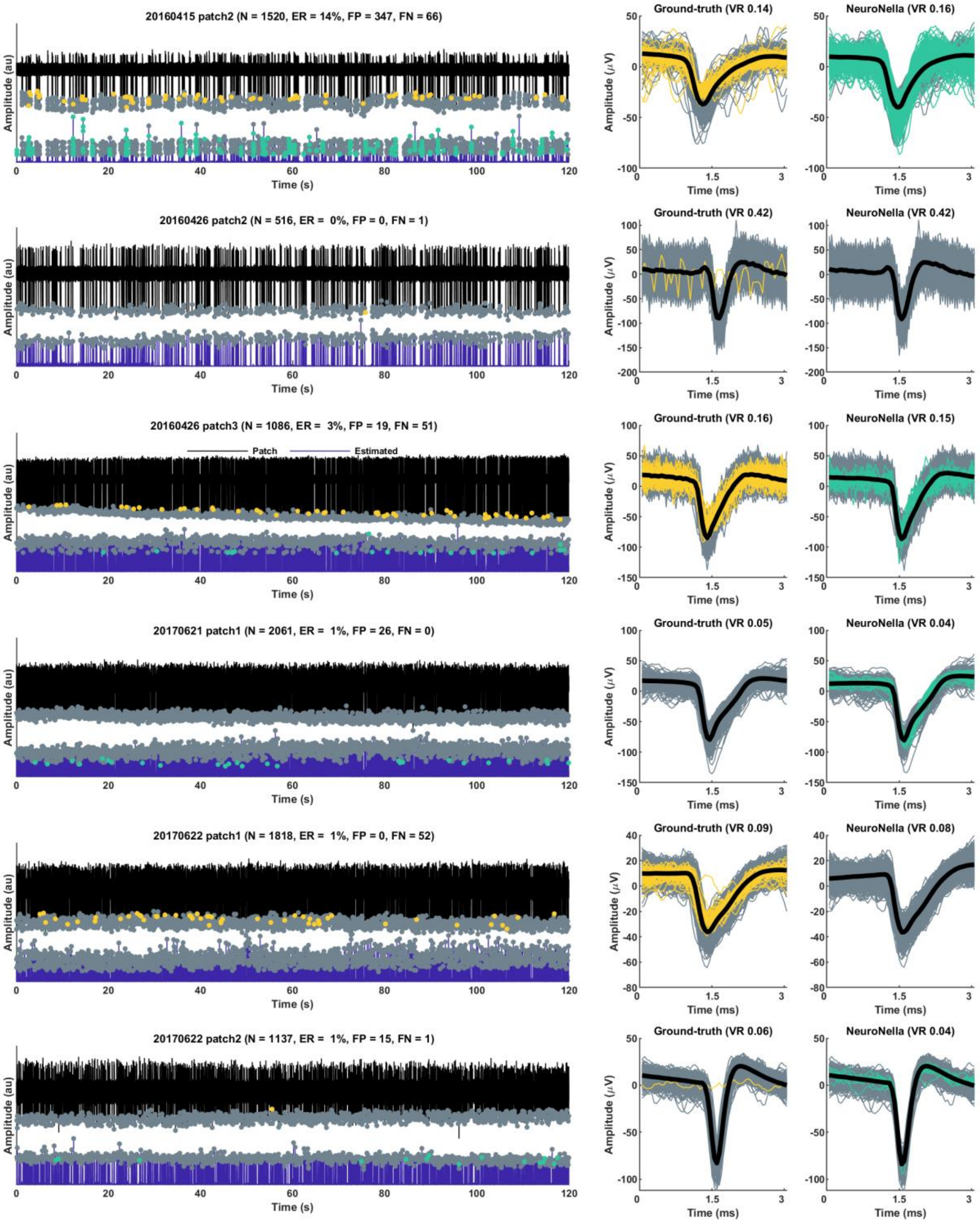

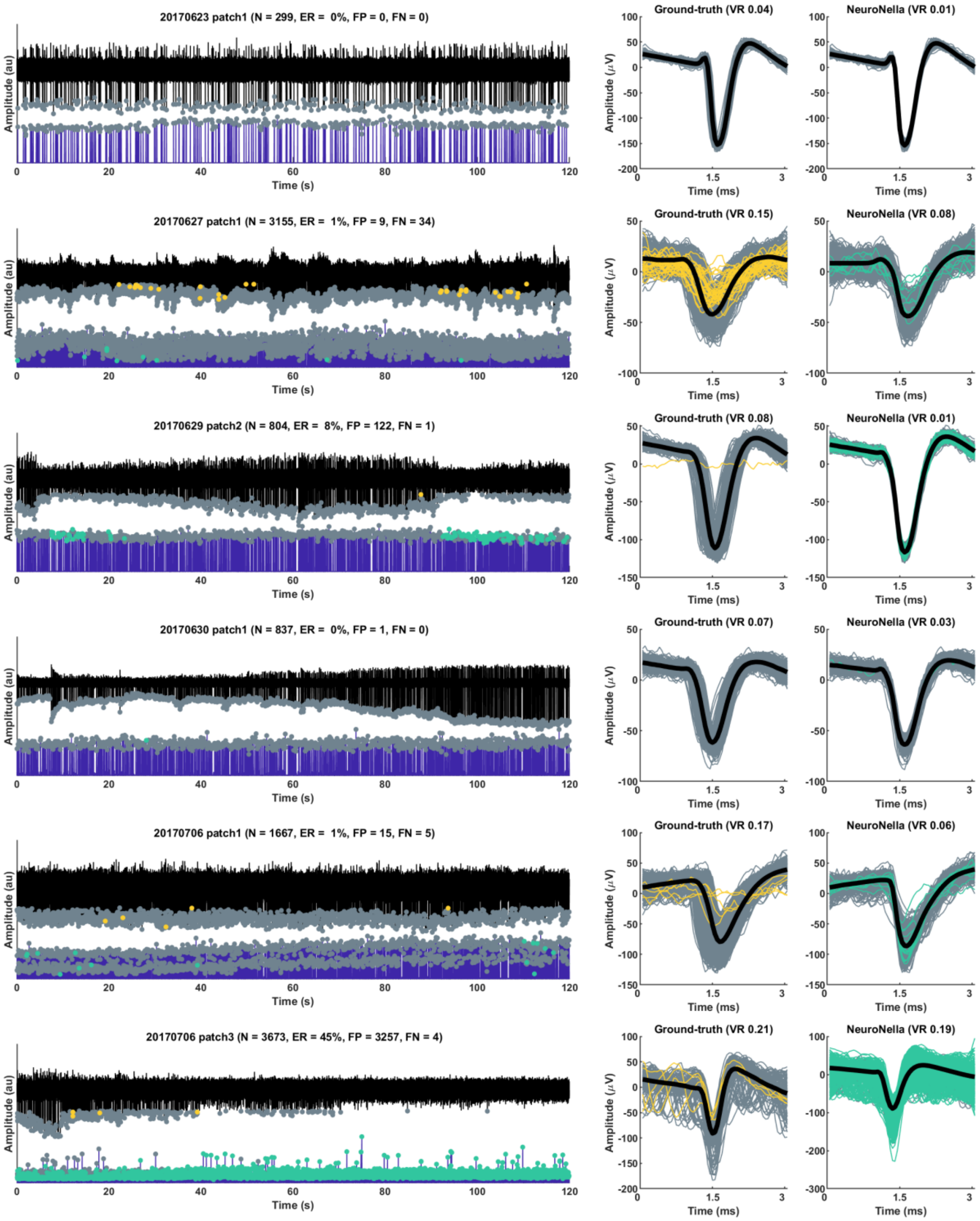

Figure A3

aaaa

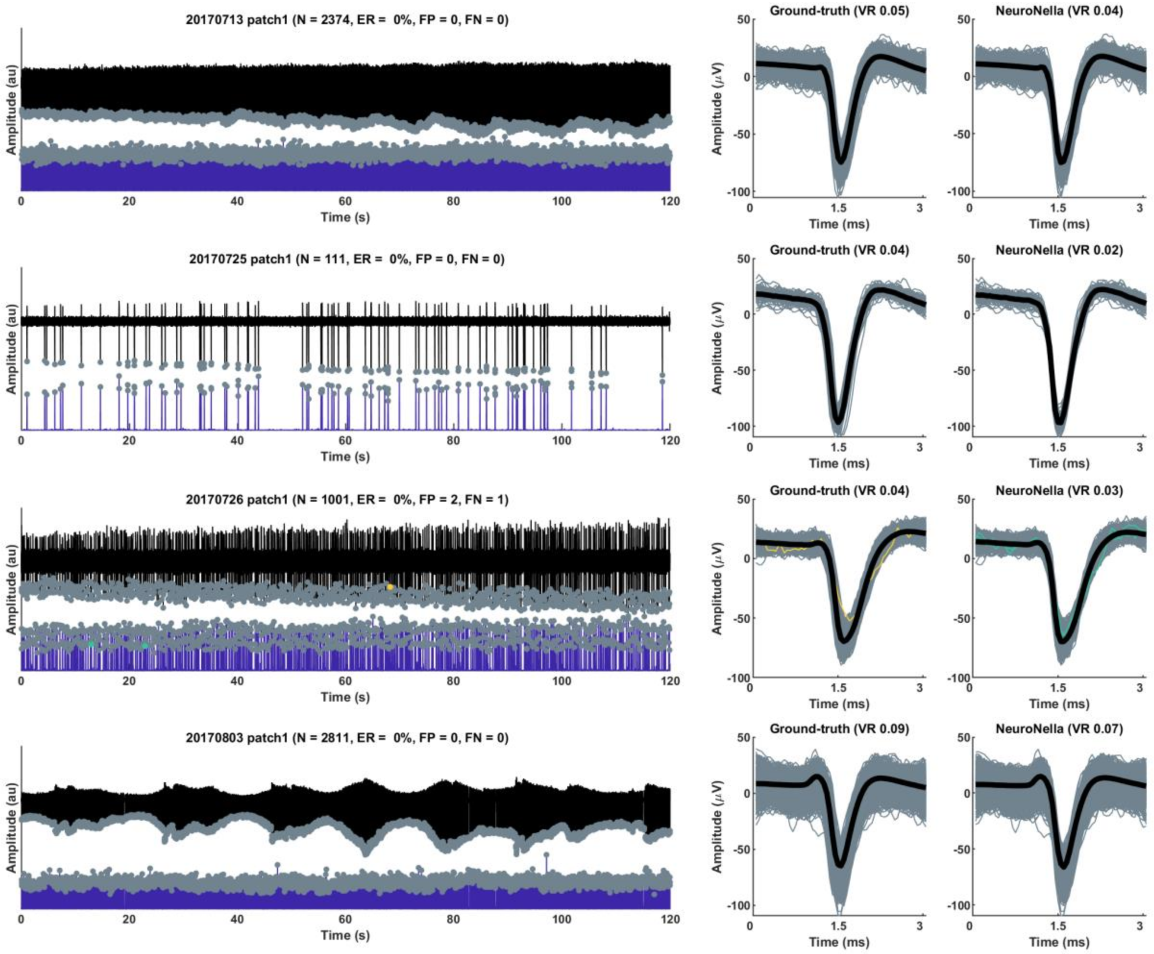
